# High-throughput Generation of Collagen Microbeads for 3D Cell Culture and Extracellular Vesicle Production

**DOI:** 10.1101/2024.11.03.621748

**Authors:** Samantha Ali, Fabiana Mastantuono, Andrea Orozco-Torres, Niloy Barua, Xin Zhou, Lizi Wu, Mei He

## Abstract

Collagen type I, a fundamental component of natural extracellular matrices across species, is an attractive material for the development of tissue engineering constructs. Nevertheless, the poor mechanical properties and thermal instability hinder its use for specific construction of 3D cultures and 3D printing. In this study, we present a microfluidic high-throughput approach for producing high-quality and uniform collagen microbeads without introducing any chemical modification. We achieved rapid and uniform collagen droplet fabrication with sizes spanning from 50 µm to 1200 µm in a production rate of up to 10000 droplets per minute. The resulting collagen microbeads can serve as numerous microbioreactors which are suspended in the culture medium without precipitation and are ideal for 3D cell growth. We demonstrated excellent cell compatibility, facilitating cell attachment and proliferation, as well as promoting extracellular vesicle secretion from collagen microbeads. This technology is facile and versatile for high throughput 3D cell culture, heterogeneous tissue modeling, and extracellular vesicle production, which is essential in drug delivery and drug screening.

## 1. Introduction

In tissue engineering, collagen type I is recognized as a promising and attractive material due to its biocompatibility and low immunogenicity[1, 2]. Collagen type I plays a critical role in providing structural integrity and supporting diverse tissue conformations, which is the primary building block of the extracellular matrix (ECM) in virtually all vertebrates and many invertebrates[3]. Collagen is able to promote ligands attachment, facilitate cell adhesion, and subsequently trigger signaling pathways that govern cell proliferation and matrix organization[4, 5]. Therefore, cells could respond to this natural ECM-derived material to initiate native biological behavior[6–8]. This is further evidenced by several FDA-approved collagen products for therapeutic and regenerative use associated with favorable clinical outcomes[9–12]. Despite of remarkable biological and regenerative properties, it has been challenging for developing collagen in 3D tissue engineering, due to its inadequate mechanical strength, low viscosity, tendency to undergo irreversible gelation, and inability to retain shear-thinning properties in 3D bioprinting[13]. In order to overcome such drawbacks, chemical modification of collagen materials has been studied, such as electrospinning with polyL-lactide [14] and chemical hybridized collagen type I [15, 16], as well as combining with different crosslink modalities including glutaraldehyde, genipin, and UV light[17]. However, the inherent natural properties of collagen could be altered, which consequently impairs the cellular biological interaction and function.

Developing collagen as a bioink for 3D bioprinting has proven challenging due to its limited mechanical properties.[18]. Some attempts utilized collagen blends to enhance material mechanical properties by leveraging various printing strategies to form collagen-based 3D constructs[19, 20]. It has been demonstrated that extrudable bio-inks can be made from granular hydrogels[21, 22]. The use of collagen and gelatin nanoparticles in granting cell binding sites to starch bio-inks has been demonstrated to create a material network with great heterogeneity ideal for replicating tissue environment[1]. Furthermore, in another study, 3D printed collagen type I supported by polyacrylamide microbeads was used to facilitate cell migration and promote microtissue formation, demonstrating a heterogeneously populated engineered tissue structure[23]. Hence, integrating collagen in the form of microbead into extrusion-based bioprinting could improve the broad applicability of collagen in tissue engineering. In this context, microgels often feature significant porosity due to the interconnected interstitial porous structures during formation [24]. This microporosity enables cells to navigate the complex structure of the material, simulating in vivo like environments, and promoting various cellular signals that stimulate endogenous cellular behavior. Particularly, 3D culture systems enhance the production of extracellular vesicles (EVs) and exhibit transcriptomic profiles similar to those found in vivo[25]. However, batch emulsion microgel generation is often polydisperse from resulting microgel contents, due to limited control over the formation of individual particles[26]. Microfluidic droplet generator systems offer the ability to fine-tune the flowing phases and control the number of cells per microgel precisely at the single-cell level[27–29]. Collagen microgels have been fabricated using microfluidic devices with great uniformity [30, 31] and used as micro-molding beads with cells grown around to facilitate migration and microtissue formation[32].

While collagen type I is frequently used in 3D tissue engineering, its direct application as an unmodified material for 3D bioprinting with live cells has not been explored in the context of 3D tissue modeling. We introduced a high-throughput droplet-based microfluidic platform for producing natural collagen type I droplets as numerous 3D microbioreactors at a large scale. By design, characterization, and optimization of microfluidic droplet fabrication, we can precisely tailor the concentration and size of collagen microbeads as building blocks for versatile downstream applications. This platform has unique features, including (i) tunable generation of collagen microbeads ranging in sizes from 45 to over 1000 µm and collagen density from 0.25 mg ml^−1^ to 3 mg ml^−1^; (ii) self-contained and biocompatible cellular encapsulation within microbeads structure for high-throughput 3D culture, (iii) seamless integration of collagen based microbeads achieving excellent rheological properties for high-fidelity 3D bioprinting; and (iv) enabling 3D biomanufacturing of extracellular vesicles at large scale. This innovative technology will provide an enabling approach for a wide range of applications in developing clinically translatable biomaterials, drug delivery and screening, tissue engineering, and regenerative medicine.

## 2. Experimental Section

### 2.1 Design and Fabrication of microfluidic device

The master mold for the microfluidic droplet generator was printed using a CADworks3D µMicrofluidics Printer using the following standard operating protocol: 50s base curing time, one base layer, layer thickness of 50 μm, 6 seconds of curing time. After printing, the master mold was carefully removed and washed with 2-propanol and dried with nitrogen to remove uncured resin. The mold was then post-cured under 390 - 410 nm UV light at 40 mW/cm^2^ for 5 minutes. The UV cured resin mold was then used to fabricate the polydimethylsiloxane (PDMS) (Sylgard) microfluidic device. An array of three microfluidic droplet generators designed in the mold using Autodesk AutoCAD software (Fig S1, supporting information). Each microfluidic device consists of two microfluidic flow-focusing channels to introduce fluorinated oil (RAN Biotechnologies) and collagen type I fluid (Advanced Biomatrix) in the cold room at 4⁰C. The collagen-in-oil droplets were formed in the channel intersect after squeezing through a 75 μm nozzle and collected at the outlet of the device. The continuous phase (oil) channels were designed to be 300 μm in width, while the dynamic phase (collagen type I) was 100 μm in width to ensure the minimum droplet size of 100 μm and the maximum droplet size of 1400 μm with variable pressure. PDMS base and curing agent were mixed at a ratio of 10:1 and mixed well for 3 minutes. The mixture was degassed in a vacuum chamber to remove air bubbles and poured on top of the master mold, then degassed again to remove any remaining air bubbles. Afterwards, the PDMS devices were cured at 75 °C for 3 hours. Then the cured PDMS was de-molded and hole-punched by a 1-mm biopsy puncher with a plunger (Integra Miltex, USA). The PDMS microfluidic device and a glass slide were then exposed to oxygen plasma for 30 seconds (Harrick Plasma Cleaner PDC-001-HP) and bonded together to complete the microfluidic device fabrication.

### 2.2 Collagen droplet generation

Initially, collagen type I from bovine skin at a concentration of 9.9 mg/ml (Advanced Biomatrix) was diluted in PBS or complete media into concentrations of 0.25, 0.5, 1, 2, 3 mg/mL. This procedure was performed in a cold room at controlled temperature of 3-5°C. Droplet generation experiments were performed inside of a portable fridge (BougeRV, Rowland Heights, CA, United States) at a temperature of 2°C. Dynamic and continuous flow phases were controlled by PreciGenome Pressure Controller to generate droplets. To investigate optimal droplet size range, both phases were subject to different pressure ratios as reported in Figure 1. To encapsulate cells within collagen, cell densities ranging from 0.5 to 25 ×10^6^ cells/mL were mixed in 1mg/mL collagen dilution. Collagen-in-oil-droplets were crosslinked and solidified at 37°C by placing into a humidified incubator with 5% CO_2_ for overnight and then washed with 1% PBS-BSA (w/v) to remove oil phase. Thereafter, collagen droplets are referred to as collagen microbeads.

**Figure 1.**
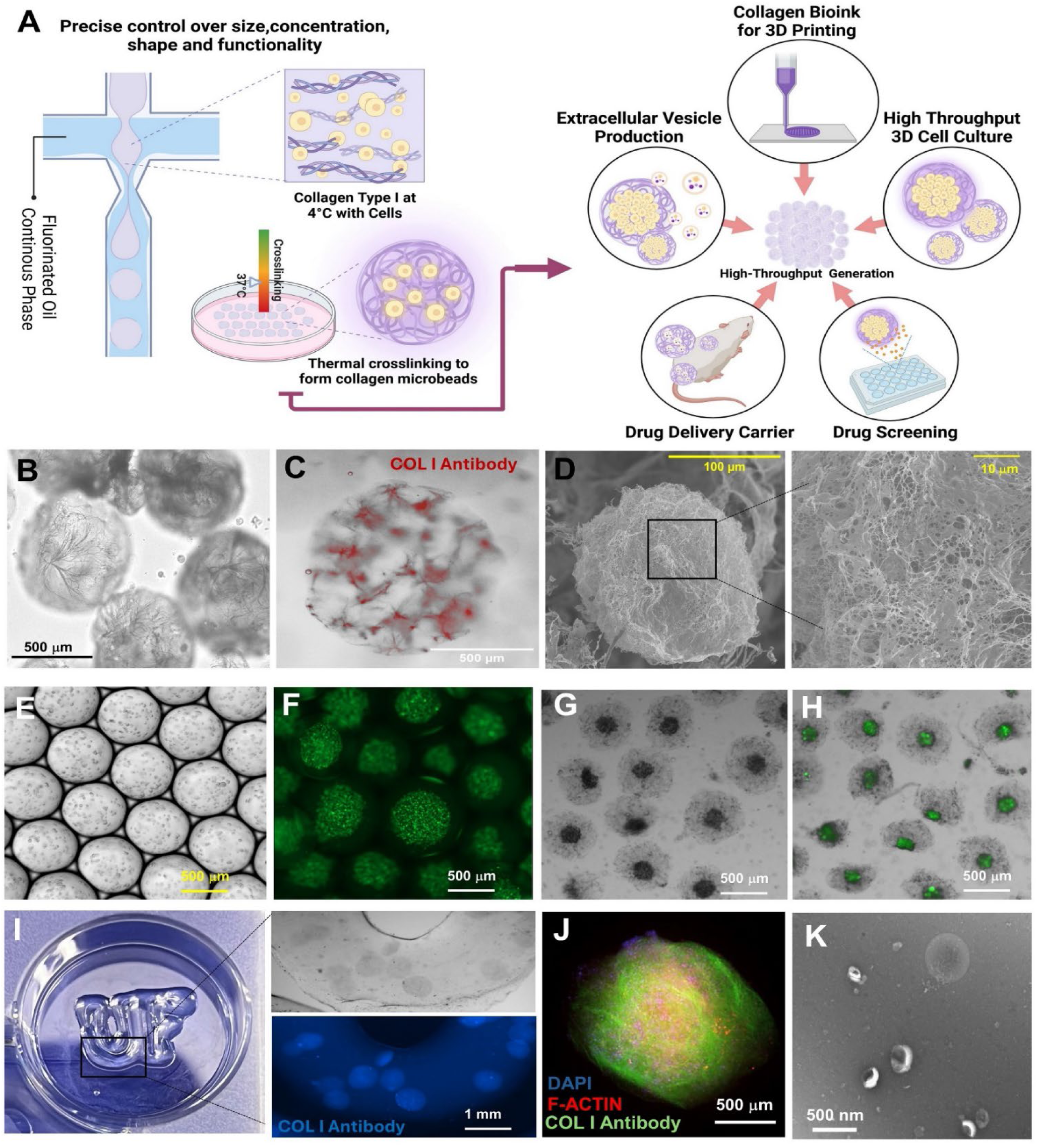
High throughput generation of collagen microbeads for versatile downstream applications. **(A)** Schematic representation of the experimental process for generating collagen microbeads in high throughput, which exhibits versatile applications in 3D bioprinting, 3D cell culture, EV production, and associated drug delivery and drug screening. **(B)** Microscopic image showing the morphology of generated collagen microbeads with typical fibrous collagen network stained in red**(C)**. **(D)** SEM image showing the collagen fibrous network under micron scale. **(E)** Microscopic image demonstrates the A549 cells encapsulated in collagen microbead droplets and produced in large scale. Cells are tagged with GFP showing green fluorescence **(F)**. Encapsulated live cells are viable and continuously growing into spheroids in Day 3 in bright field image **(G)** and fluorescence microscopic image **(H)**. **(I)** Picture of 3D printed five-layer constructs using collagen microbeads encapsulated with A549 cells as the bio-ink. The zoom in picture showed the live cell microbeads uniformly distributed in the 3D printed constructs. **(J)** Confocal image showing the spheroid growth within collagen microbioreactor which exhibited optimal cell attachment and stretch into a tissue-like structure. **(K)** TEM image demonstrated the EV production harvested from 3D spheroid culture in collagen microbioreactors.

### 2.3 Cell culture

Human lung cell lines were utilized for the experiments, including H322 bronchioalveolar carcinoma cells, A549 adenocarcinoma alveolar basal epithelial cells, and BEAS-2B non-tumorigenic lung epithelial cells, which were all tagged with GFP. The H322 and A549 cell lines were cultured in Dulbecco’s Modified Eagle Medium (DMEM) supplemented with 10% heat-inactivated fetal bovine serum (FBS) and 1% antibiotics-antimycotics. The BEAS-2B cell line was maintained in Roswell Park Memorial Institute 1640 Medium (RPMI 1640), similarly supplemented. Upon reaching desired confluence, cells were washed with PBS and detached using 0.25% Trypsin-EDTA (Thermo Fisher Scientific). Following a 5-10 minute incubation, cells were centrifuged at 300g for 5 minutes to pellet. Cell counting was conducted to determine the number of cells per droplet. Subsequently, live cells were suspended in collagen type I and processed through the microfluidic platform for encapsulation into collagen microspheres. The cell-laden microspheres were incubated at 37°C in a humidified incubator with 5% CO_2_ to promote cell proliferation and function. Cell viability in collagen microbioreactors was measured by performing resazurin based assay PrestoBlue HS every day for 7 days.

### 2.4 Collagen microbead bio-ink fabrication, characterization and 3D printing

To prepare the collagen microbead-enriched bio-ink, 4% gelatin type A and 3.25% sodium alginate (AG gel) were dissolved in PBS at 60°C for one hour, using a magnetic stirrer set to 600 rpm. Once the hydrogel solution was fully dissolved, the temperature was reduced to 37°C and collagen microbeads were added into hydrogel solution. Different concentrations of collagen microbeads relative to AG gel (v/w%) were evaluated, including 0.1:1, 1:2, 1:1, and 2:1. The prepared bio-ink was loaded into a sterile 30 cc cartridge and extruded through a 25G tapered tip needle with an internal diameter of 260 μm. The printing process was performed using EnvisionTEC Bioplotter, which utilized a temperature-controlled cartridge and nozzle, with the printing temperature maintained at 27°C. A five-layer construct measuring 12 x 12 x 1 mm was printed using an extrusion flux of 8 mm/s and a pressure of 2 Pa. The printing chamber temperature was kept between 21-23°C to facilitate immediate gelatin crosslinking. Post-printing, the scaffolds were immersed in 100 mM calcium chloride for one minute to achieve alginate crosslinking, thereby ensuring the integrity of the 3D structure. Fluorescent images were obtained by conjugating a collagen type I antibody with Alexa Fluor 405.

### 2.5 Rheology Measurement

The rheological properties of the collagen microbead bio-inks were assessed using an MCR 702 rheometer (Anton Paar) equipped with a 25 mm diameter parallel plate geometry. The experiments were conducted at a controlled temperature of 25°C. Initially, strain sweep tests were performed to determine the linear viscoelastic region. Subsequent measurements were carried out within this region. Oscillatory frequencies were measured in the linear viscoelastic range by performing oscillatory frequency sweeps from 15 Hz to 10^3^ Hz at a strain amplitude of 1%. Complex viscosity was measured as a function of angular frequency to evaluate the viscoelastic behavior of the collagen bio-ink. The viscoelastic properties were compared across different concentrations of collagen microbead to AG gel ratios, including 0.1:1, 1:2, 1:1, and 2:1, alongside control samples of pure AG gel and alginate.

### 2.6 Extracellular vesicle isolation and characterization

For harvesting extracellular vesicles from 3D spheroids cultured in collagen microbioreactors, the mouse myoblast cell line C2C12 was used with a density of 10×10⁶ cells and resuspended in 1 mL of 1 mg/mL collagen type I to form 300-μm collagen cell microbeads, resulting in approximately ~141 cells per microbead. The resulting microbioreactor cultures were grown for 3 days in DMEM supplemented with 10% FBS, after which the media was switched to DMEM with 10% exosome depleted media and grown for an additional 48 hours before harvesting extracellular vesicles. The collected media was centrifuged at 300 g, 2000 g, and 10,000 g for 5, 20, and 30 minutes respectively to remove cellular debris, then transferred to an ultracentrifuge tube. A 3 mL 30% (w/v) sucrose solution was placed at the bottom of the ultracentrifuge tube, which was then centrifuged at 100,000 g for 90 minutes at 4°C. The sucrose cushion was subsequently transferred to a new ultracentrifuge tube, 10 mL of ice-cold PBS was added, and the mixture was gently pipetted to disrupt the sucrose cushion. Another ultracentrifugation at 100,000 g for 90 minutes at 4°C was followed afterwards. Finally, the supernatant was removed, and the pellet was resuspended in 400 μL of PBS to collect EVs.

Nanoparticle tracking analysis was performed using the ZetaView system (QUATT, Particle Metrix Inc, USA). The laser was set to a wavelength of 488 nm, with a sensitivity set to 80 and a shutter time of 100. Measurements were taken over 90 seconds at the highest video resolution across all 11 positions.

Total protein in EVs was quantified using the Pierce BCA Protein Assay. The 10 µl of isolated EVs were lysed with 1 µl of cold RIPA buffer, incubated on ice for 10 minutes, and then sonicated for 30 seconds. According to the vendor’s protocol, reagents A and B were mixed in a 24:1 ratio to prepare the working buffer. In a separate tube, 10 µL of the working reagent was added to the lysed EV sample, followed by incubation at 60°C for 1 hour. After cooling to room temperature, the spectrum absorbance was measured at 562 nm.

Total RNA was extracted from isolated EVs using the miRNeasy Mini Kit following the manufacturer’s instructions. EVs were lysed with QIAzol and incubated for 5 minutes at room temperature. Chloroform was added, mixed vigorously, and incubated for 3 minutes at room temperature, followed by centrifugation at 12,000x RCF for 15 minutes at 4°C to allow phase separation. The aqueous phase was extracted, mixed with 100% ethanol, and transferred to the RNeasy Mini column. The column was centrifuged at 8,000xg for 30 seconds, washed with 700 µl of reagent RWT, and centrifuged again at 8,000×g for 30 seconds. Next, 500 µL of reagent RPE was added and centrifuged for 2 minutes at 8,000×g. The membrane was dried centrifuging at 8000×g speed for 1 minute. RNA was eluted by adding 30 µL of RNase-free water to the column and centrifuging at 8,000×g for 1 minute, with a repeat application of the eluate. RNA quality was assessed using the 260/280 nm ratio, and sample concentration was analyzed using total RNA 6000 Pico Kit according to the manufacturer’s protocol.

### 2.7 Image acquisition and analysis

Agilent Cytation 5 Cell Imaging Multi-Mode Reader equipped with an inverted fluorescent microscope (widefield) was used for transmitted light and fluorescent cell and microbead imaging. Magnifications were achieved using 4× and 20× objectives. Images were processed using Gen5 software. Collagen droplet and cell spheroid sizes were quantified using primary cellular analysis capabilities of Gen5 software using transmitted light and GFP imaging channels. Postprocessing of fluorescent imaging includes adjusting contrast and brightness to denote cell structure within spheroids and 3D scaffold.

### 2.8 Scanning Electron Microscope

Collagen microbead samples were rinsed in distilled water three times, frozen at −80°C and lyophilized (Labcono 2.5 L) overnight. Hitachi Scanning Electron Microscope (SEM) to visualize and probing fibril structures within collagen microbeads and collagen microbead-hydrogel bio-inks (University of Florida ICBR Electron Microscopy Core Facility, RRID:SCR_019146). Aperture and stigmata corrections were done before sample images were obtained. ImageJ was used to analyze the microstructures of the samples.

### 2.9 Confocal Microscopy

Mouse myoblast C2C12 cells were encapsulated in type I collagen and cultured for 14 days. Following this period, the cells were fixed in 4% paraformaldehyde, permeabilized with 0.01% Triton X-100, and blocked with FBS as described [33]. Resulting 3D microtissues were then stained for F-actin using Phalloidin AlexaFlour594, Collagen type I antibody conjugated to Cy3, and Hoechst for nuclei and imaged in a Leica Stellaris 8 WLL Spectral Confocal Microscope (University of Florida ICBR Cytometry Core Facility, RRID:SCR_019119)

### 2.10 Transmission Electron Microscope

Ultrathin copper grids underwent a 1-minute glow discharge treatment prior to use. Subsequently, 10 μL of EV samples were each applied to the treated grids and allowed them to sit undisturbed for 5 minutes at room temperature. The grids were rinsed once with distilled water and then negatively stained with filtered 2% aqueous uranyl acetate for 2 minutes. After air drying at room temperature, the grids were examined using transmission electron microscopy (TALOS TEM L120C G2 120 kV, University of Florida ICBR Electron Microscopy Core Facility, RRID:SCR_019146) to verify the morphology of EVs.

### 2.11 Statistical analysis

All results are expressed as the mean ± standard deviation (SD). Statistical analyses were done using Prism Graphpad 6.0. Statistical significance was considered when p≤ 0.05 and greatly significant when p≤ 0.001. Three independent trials were carried out unless otherwise stated.

## 3. Results and discussion

### 3.1 High throughput generation of collagen microbeads

We used a microfluidic water-in-oil approach within a flow-focusing device made from PDMS to produce monodisperse collagen microbeads (Figure 1, Figure S1 supporting information). A schematic concept illustration and potential downstream diverse applications, including 3D bioprinting, high throughput 3D cell culture, EV production, drug delivery and drug screening, are shown in Figure 1A. The collagen droplets were formed by the pinching motion of the oil phase (FC-40 or HFE-7500) and aqueous collagen phase at 4⁰C, and stabilized by the coalescence of these two immiscible phases at 37⁰C. Figure 1 B and C displayed the microscopic morphology of formed collagen droplets, showing fibrous microporous collagen networks. In Figure 1D, the SEM imaging further confirmed the typical fibrous collagen structure. Figure 1 E and F demonstrated the live cell encapsulation in uniform collagen droplets in large scale production, as well as the large-scale 3D cell culture to continue to grow as spheroids (Figure 1 G and H). Figure 1 I demonstrated the 3D printing using generated collagen microbeads encapsulated with adenocarcinomic human alveolar basal epithelial live cells (A549). The collagen microbeads with A549 cells were uniformly distributed in the 3D printed constructs (Figure 1I inserts). Figure 1J confocal microscopic imaging demonstrated the 3D cell culture within individual collagen droplet as the microbioreactor, which exhibited optimal cell growth with attaching and stretching into collagen fibrous network. Figure 1K TEM imaging is a demonstration of harvested EVs produced from 3D cell culture in collagen microbioreactors. Overall, high throughput generation of collagen microbeads provided a versatile platform for 3D bioprinting, 3D cell culture, and associated EV production, which is essential for developing drug delivery biomaterials and high throughput drug screening.

In the microfluidic droplet generator, due to the nature of the viscosities of each flow phase (e.g.,FC-40 2 cSt, HFE 0.81 cSt, collagen type I approximately 8 cSt) and channel sizes, the size distribution of collagen droplets can be controlled by regulating the pressure rate of each flow. Figure 2 investigated the relationship between the size distribution of collagen microbeads and the experimental condition on pressure ratios of the oil-to-collagen phase. We studied both FC-40 and HFE-7500 oils to investigate the formed collagen microbeads and achieved size range from 45±2 μm to 1300±200 μm, indicating a tunability to precisely control the size of collagen microbeads (Figure 2 A and D). The size distribution of droplets is proportional to oil viscosity, allowing to approximate the necessary pressure ratios for droplet formation. Figure 2B and E illustrate the production rate in terms of frequency over different sizes of microbeads distributed for each pressure ratio and oil type (B: FC-40; E: HFE-7500). The collagen droplet production rate determines the throughput and production capacity, which is ranging from ~ 6×10^3^ microbeads per minute for FC-40 oil (Figure 2C) and 1×10^4^ microbeads per minute for HFE7500 oil (Figure 2F). HFE7500 oil enables a much higher production rate (supporting information, video S1), due to the lower viscosity for efficient flow and good stability. Overall, our approach results in physically monodisperse microbeads with large scale production capability (Figure 2G). The collected collagen microbeads were incubated at 37°C overnight to stabilize their spherical shape via thermal crosslinking, which promoted non-covalent fibril assembly via primarily hydrogen bonding to form a dense fibril network for ideal 3D cell growth. After stabilization, the microbeads were washed in PBS multiple times to remove oil residues. These collagen microbeads can be stored in a PBS solution at 4⁰C for more than three months for later usage. This highly scalable process is readily available for high throughput drug screening using 3D cultured cells or serving as the bio-ink for 3D printing.

**Figure 2.**
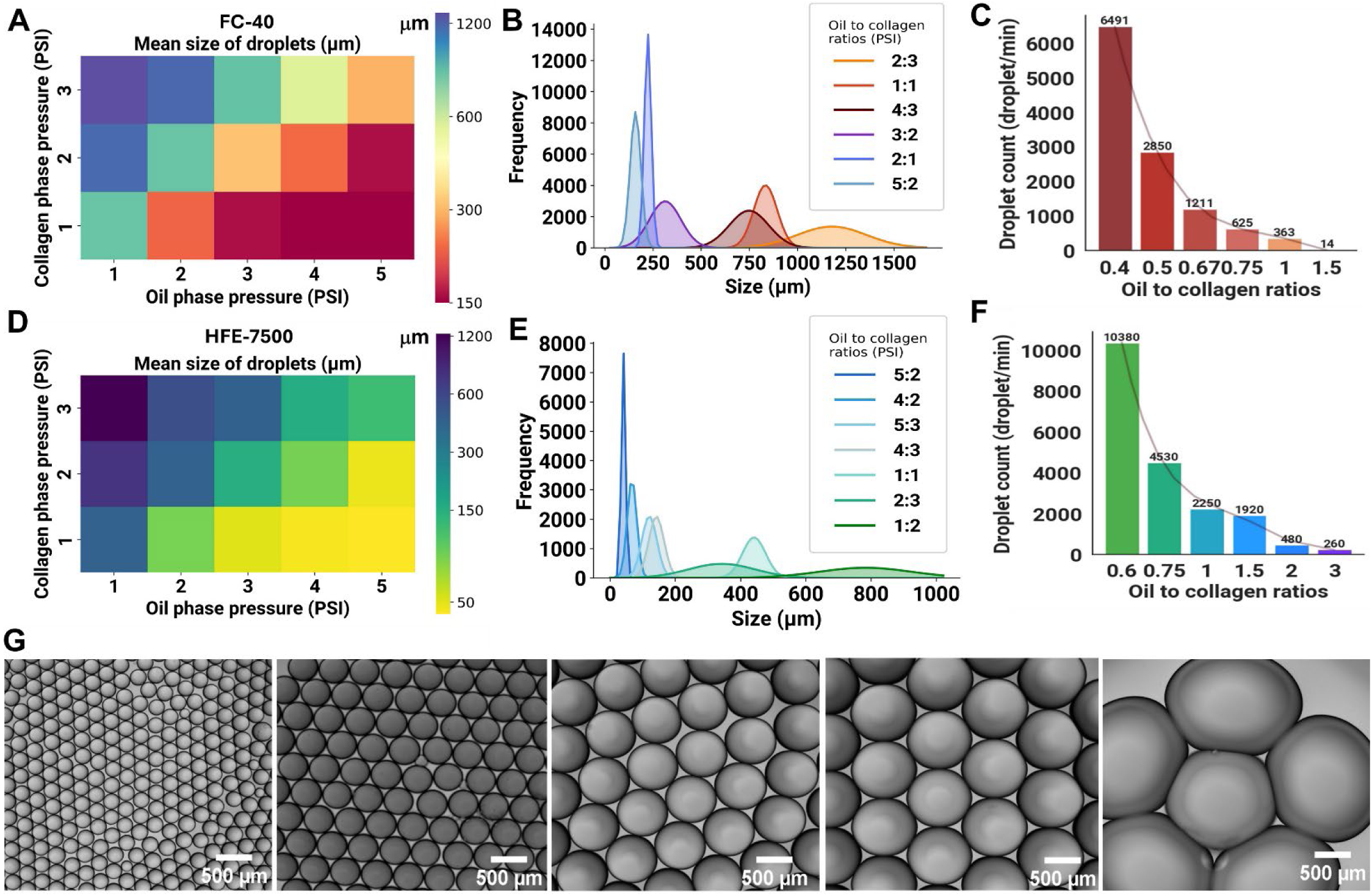
Size controlled production of collagen microbeads. **(A)** Mean droplet size distribution across various oil-to-collagen ratios using FC-40 oil. **(B)** Frequency distribution over various droplet sizes under different oil-to-collagen ratios using FC-40 oil. **(C)** Collagen microbeads production rate per minute across different oil-to-collagen ratios using FC-40 oil. **(D)** Mean droplet size distribution across various oil-to-collagen ratios using HFE-7500 oil. **(E)** Frequency distribution over various droplet sizes under different oil-to-collagen ratios using HFE-7500 oil. **(F)** Collagen microbeads production rate per minute across different oil-to-collagen ratios using HFE-7500 oil, indicating high efficiency and throughput. **(G)** Uniformity of collagen microbeads produced with 1 mg/mL collagen type I at different sizes, demonstrating the precision and replicability from the production process.

### 3.2 Characterization of microstructure of collagen microbeads

Interestingly, we also observed that 1 mg/mL collagen microbeads remained in suspension in the culture medium without precipitation, which is ideal for maintaining 3D cell growth (supporting information Figure S2, video S2). Such suspension ability could be attributed to their microstructural porosity and fibrous network. Since collagen forms the molecular mesh of 90% of all human tissue for modulating cellular behavior [34, 35], the interconnectivity and microstructure could play a vital role in the architecture of collagen microbeads. Herein, to explore the porosity and fiber density under the influence of collagen concentrations, microbeads were formed in collagen concentrations ranging from 0.25 – 3 mg/mL (Figure 3A). The resulting microstructure was observed by cryo-SEM imaging shown in Figure 3A bottom panel, which clearly illustrated the increased density of fiber network along with higher collagen concentration. For microbeads in 0.25 mg/mL collagen concentration, a slight deformation of the initial spherical shape was noted, which could be due to poor mechanical strength from low collagen concentration for crosslinking. In contrast, for concentration of 3 mg/mL, a nearly hexagonal shape was observed, suggesting an ultra-dense packing of collagen fibers within the microbead. The freeze-dried collagen microbeads (Figure 3A bottom) revealed that fibril formation is proportionally dependent on the initial collagen concentration, leading to more dense interconnected porous networks (Figure 3B) and smaller pore size (Figure 3C). The thicker fibers were observed as collagen concentrations increase (Figure 3D), which is responsible for providing tensile strength and viscoelasticity to tissues, indicating that denser microbead structures may offer greater mechanical robustness and reduced elasticity.

**Figure 3.**
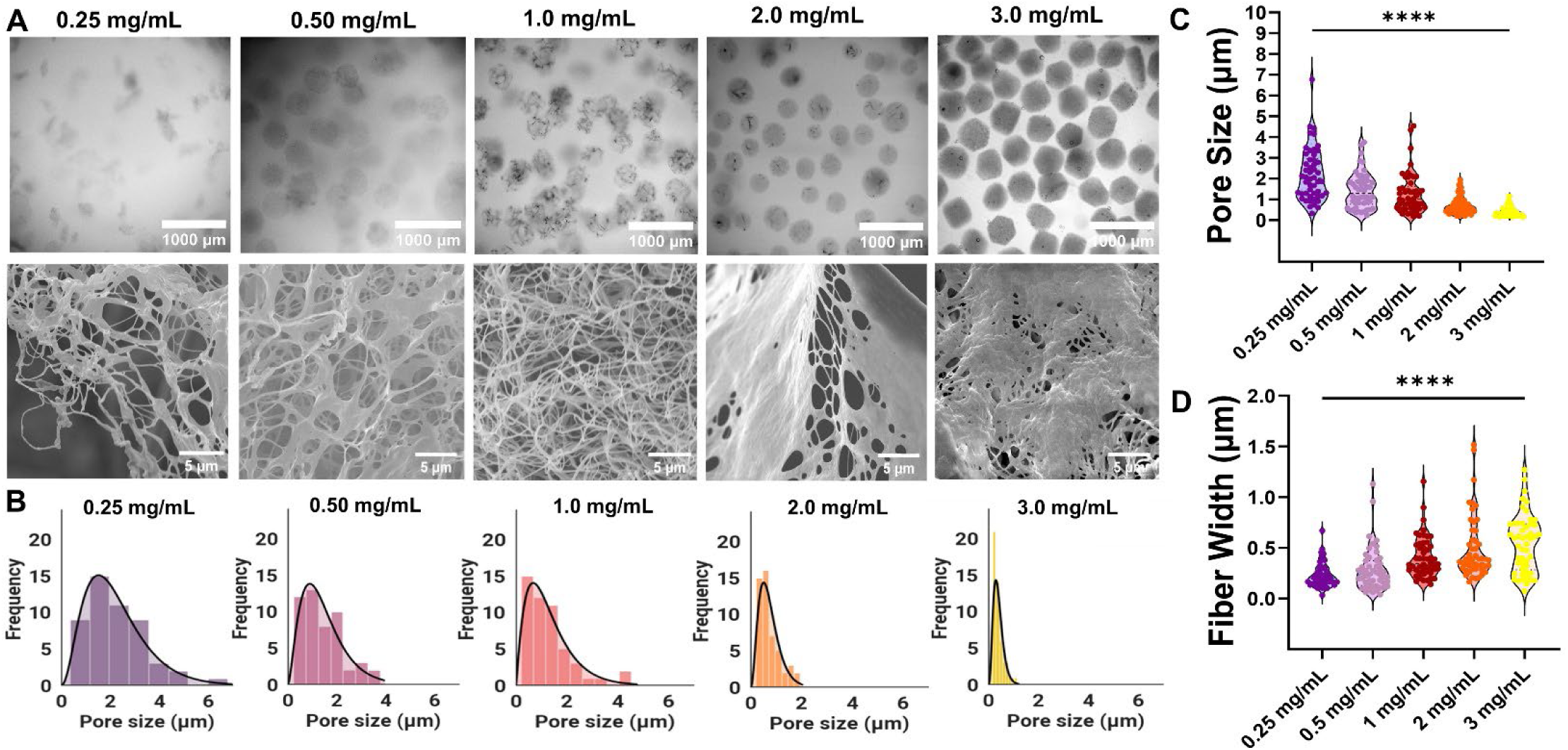
Characterization of microstructure of collagen microbeads. **(A)** Top row: Light microscopy images showing the overall microbead structure. Microbead concentrations: 0.25 mg/mL, 0.50 mg/mL, 1.0 mg/mL, 2.0 mg/mL, and 3.0 mg/mL, respectively. Bottom row: Cryo-SEM images depicting the detailed network of collagen fibrils. **(B)** Pore size distribution histograms for collagen microbeads at concentrations of 0.25 mg/mL, 0.50 mg/mL, 1.0 mg/mL, 2.0 mg/mL, and 3.0 mg/mL, respectively. **(C)** Violin plots showing the microbead pore size distribution across different collagen concentrations, highlighting a significant decrease in pore size with increasing concentration (**** p < 0.0001). **(D)** Violin plots illustrating the fiber width distribution across different collagen concentrations, indicating an increase in fiber width with higher concentrations (**** p < 0.0001).

Given that microbead suspensions exhibit unique viscoelastic properties beneficial for extrusion-based 3D printing, specifically shear thinning where the solution’s viscosity decreases as force and shear rate increase[21, 24, 26]. Therefore, a rheological evaluation was performed shown in Figure 4. Approximately, 5×10^4^ of collagen microbeads were mixed gently in alginate-gelatin (AG) in a v/v proportion. Viscosity measurements indicate that the lowest fraction of microbead included in the AG hydrogel bio-ink has slight effect on the viscosity over time and angular frequency, respectively (Figure 4A and B). To further explore the shear-thinning behavior, the storage modulus (G’) and the loss modulus (G’’) both over the time and angular frequency were measured in Figures 4C and D, respectively. The result reveals that ratios between collagen microbead and AG gel not only controlled their elasticity but also exhibited increased mechanical robustness as the collagen content was increased. Particularly for all formulations (0:1, 1:2, 1:1, 2:1), G’ remains relatively stable over time, indicating that the elastic properties of the bio-inks do not significantly change within the measured timeframe. The content of collagen microbeads maintained suitable shear-thinning behavior for continuous filament deposition and shape fidelity in 3D printing. The loss modulus (G’’) analyses indicated increased elasticity at higher deformation rates with higher content of collagen microbeads (Figure 4E).

**Figure 4.**
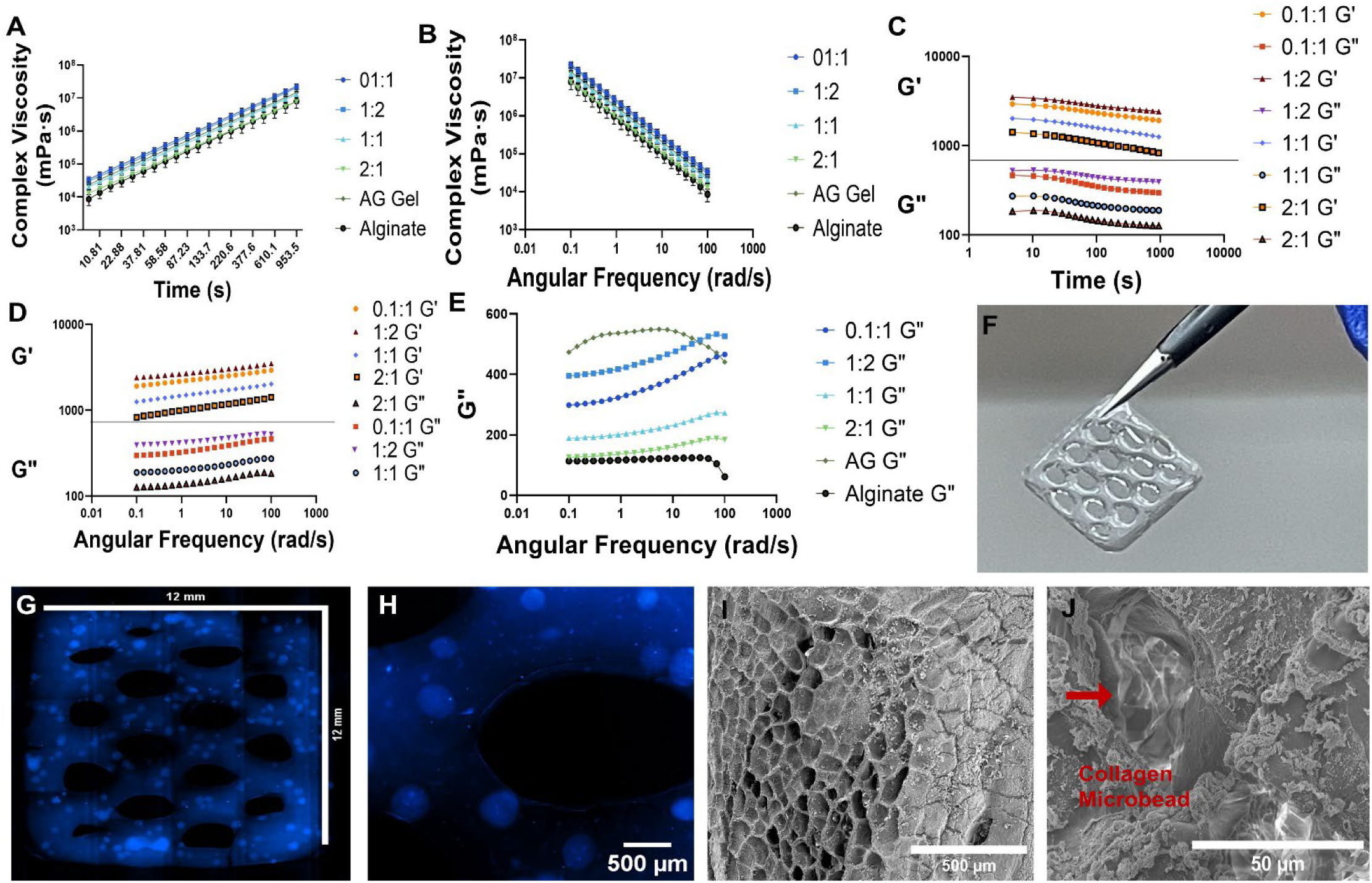
Rheological characterization of collagen microbeads for 3D printing. **(A)** Complex viscosity as a function of time for collagen-alginate gelatin bioink at different composition ratios (0:1, 1:2, 1:1, 2:1, and pure alginate), alongside an alginate gel control, demonstrating the shear-thinning behavior over time. **(B)** Complex viscosity versus angular frequency for the collagen-alginate gelatin bioink formulations, highlighting the frequency-dependent viscosity characteristics. **(C)** Storage and loss moduli as a function of time, indicating the viscoelastic properties of the bioink with varying collagen-alginate ratios. **(D)** Storage and loss moduli versus angular frequency, showcasing the stability of the viscoelastic properties across a wide range of frequencies. **(E)** Loss tangent as a function of angular frequency for the bioink formulation, illustrating the transition between predominantly viscous to predominantly elastic behavior. **(F)** Picture of 3D printed honeycomb mesh constructs. **(G)** Fluorescence microscopic image of 3D printed honeycomb mesh constructs with collagen labeled in blue (collagen type I antibody-AlexaFlour 405). **(H)** Enlarged honeycomb mesh constructs showing the uniform distribution of collagen microbeads after 3D printing (collagen type I antibody-AlexaFlour 405 in blue). **(I)** SEM image showing the overall 3D printed honeycomb mesh constructs. **(J)** Enlarged honeycomb mesh constructs after 3D printing under SEM imaging. Red arrow indicates the collagen microbead.

To proof of concept using collagen microbeads as the bio-ink, we used a 1:1 ratio of collagen microbeads to AG gel, and printed through a 21-gauge nozzle (508 μm). To be able to image the distribution of collagen microbeads in 3D printed constructs, we stained collagen microbeads with collagen type I antibody-AlexaFlour 405. We used optimized printing parameters to achieve a multi-layered mesh 3D construct measuring 12×12 mm^2^ (printing speed of 6 mm/s and a pressure of 1.2 PA) (Figure 4F). Our observations confirmed that collagen microbeads were extruded smoothly and evenly within the AG precursor hydrogel (Figure 4G and H). The bridging segments between two perpendicular sections of the filaments displayed the microbeads adjacent layers as characterized under microscopy (Figure 4G). Due to the thermoplastic behavior of gelatin, the printing temperature was controlled around 25 to 30°C to enable shape-controlled printing. We examined the microstructure of 1:1 AG-Collagen microbeads using SEM. Figure 4I highlights the highly porous structure of the overall 3D printed constructs, which exhibited a network resembling of honeycomb-like pattern with an average pore size of 106.09 μm ± 34.8 (Figure S3). In the inner hierarchal structure, we can observe fibrous structures of collagen microbeads (Figure 4J).

### 3.3 Collagen microbeads for high throughput 3D Cell culture

In the microfluidic system, the encapsulation of cells in droplets is influenced by the random spatial distribution and the timing of cell arrival at the droplet formation site. This randomness adheres to the statistical principles of Poisson distribution as shown in equation 1. The variable *P* represents the fraction of droplets expected to contain a quantity of *x* cells. The parameter *λ* denotes the mean number of cells per individual droplet and is computed by multiplying the cell concentration and the volume of each droplet. The query was set for cell concentrations ranging from 1×10^6^-20×10^6^ cells/mL and microbead sizes of 100-300 µm.

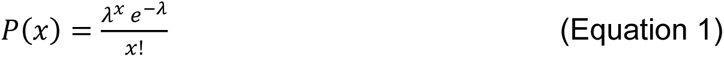

We demonstrated that our microbead cell encapsulation follows a Poisson distribution along collagen microbead size ranging from 100 μm to 300 μm in Figure 5 A-C. The higher cell concentration and the larger microbead size, the closer distribution resembled with the Gaussian distribution, which ensures a more uniform distribution of cells within each microbead. This uniformity is essential for applications requiring consistent cell numbers, such as constructing 3D organoids and microtissue models. On the other hand, in 3D culture systems where counting cells to determine growth is challenging. This approach provides a reliable baseline for estimating cell proliferation. For instance, we utilized the H322 human adenocarcinoma cell line and the BEAS-2B human epithelial alveolar cell line, chosen for their average diameters of 12 µm and 24 µm, respectively. As shown in Figure 5D and Supporting Information Video S3, we observed higher cell density encapsulation within collagen microbeads along with increased cell concentrations. We compared the observed mean encapsulation for cell concentrations of 1×10^6^, 2.5×10^6^, and 5×10^6^ to the theoretical cell encapsulation by the Poisson distribution as shown in Figure 5E, which indicates a good correlation (R^2^=0.98). By applying the Poisson distribution, we can approximate cell encapsulation within microbeads, providing a reliable means for quantifying cell densities within collagen microbeads for 3D culture.

**Figure 5.**
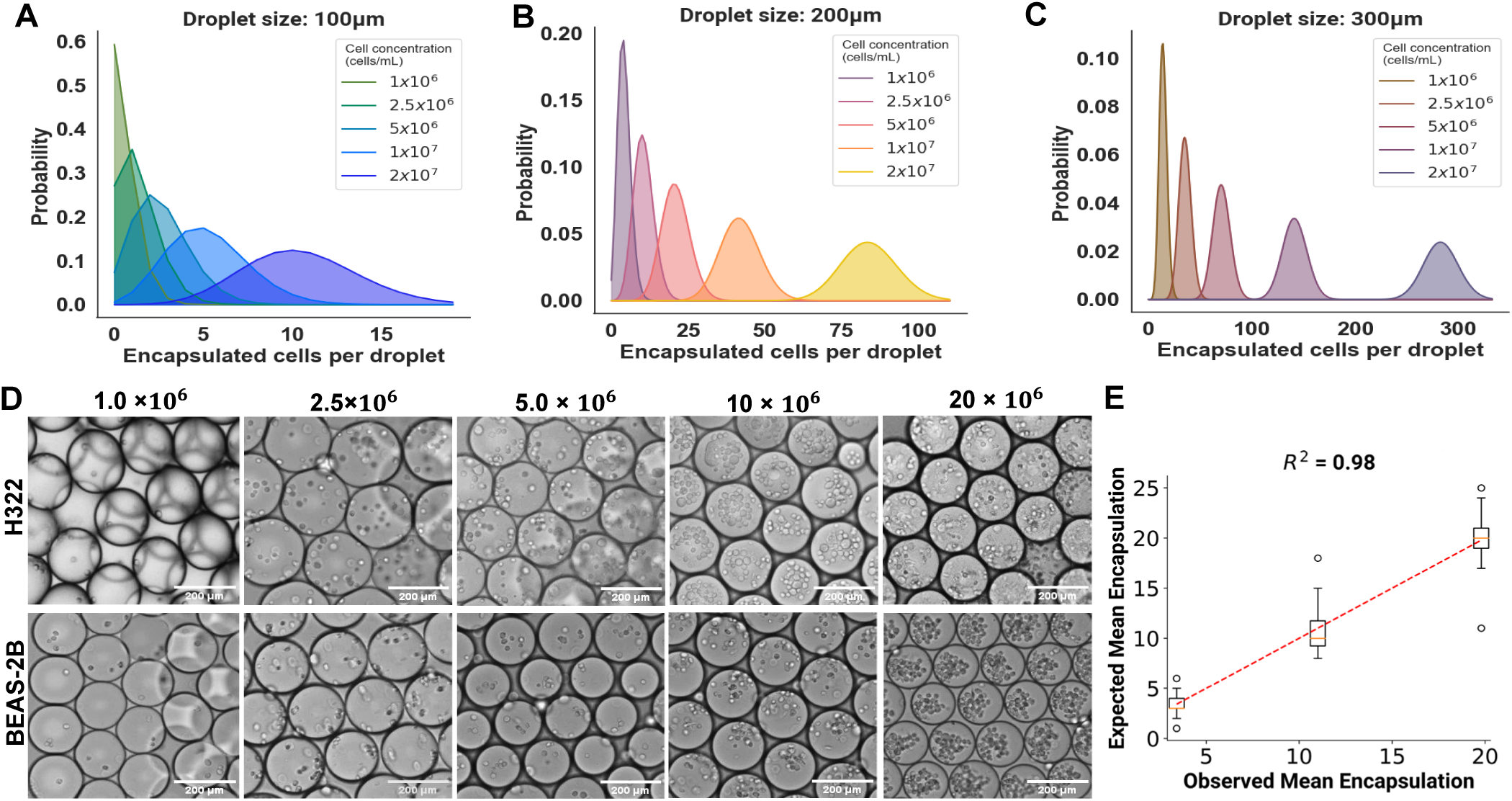
High precision collagen microbead cell encapsulation. **(A)** Cell encapsulation probability per collagen microbead with calculated λ for microbead sizes ranging from 100 μm, 200 μm **(B)**, to 300 μm **(C)**. **(D)** Representative bright field (BF) images showcasing the efficiency of H322 and Beas2B cells encapsulated in collagen microbeads across various cell densities. Cell densities are the 1×10^6^, 2.5×10^6^, and 5×10^6^ cells per mL. Scale bars represent 200 µm. **(E)** Correlation of the experimental value (black data points) of cell encapsulation per microbead with predicted values from Poisson distribution (red dash line).

For testing the suitability of collagen microbead as microbioreactors in 3D cell culture, we tracked the growth of both A549 human lung carcinoma epithelial cells and BEAS-2B human epithelial alveolar cells within our collagen microbeads (Figure 6). A 1 mg/mL collagen solution was diluted in a complete growth medium and combined with a cell density of 5×10^6^ cells/mL. This mixture was processed through our microfluidic platform to produce collagen microbeads of approximately 200 µm in diameter and each contained approximately 21 cells (Figure 6A and D). The microbeads were incubated at 37°C overnight to facilitate collagen thermal crosslinking and cell growth was evaluated for 7 days to investigate cell viability (Figure 6B, 6E). We measured the spheroid size over a period of 7 days (Figure 6C, 6F) and performed a resazurin-based viability assay to monitor cell growth within the spheroids. This assay allowed us to assess cell proliferation and viability over time quantitatively (Figure 6G, 6H). The distinct spheroid morphology from two different cell types was observed through the days of growth. At Day 3, the fibrous structure of collagen can no longer be observed in A549 spheroids through our widefield microscopic images. For BEAS-2B cells at the same time point, the fibrous structure of collagen is still distinctive. We suspect that fibrous structure observed in BEAS-2B cell growth could be due to matrix deposition, because epithelial cells tend to remodel their ECM [36, 37]. Overall, both cell lines remained viable and capable of forming spheroids within the collagen microbeads for more than a week. The A549 cells proliferated more rapidly and formed larger spheroids than BEAS-2B cells, which indicates the versatility of collagen microbeads in supporting various cell types propelling their unique growth profiles.

**Figure 6.**
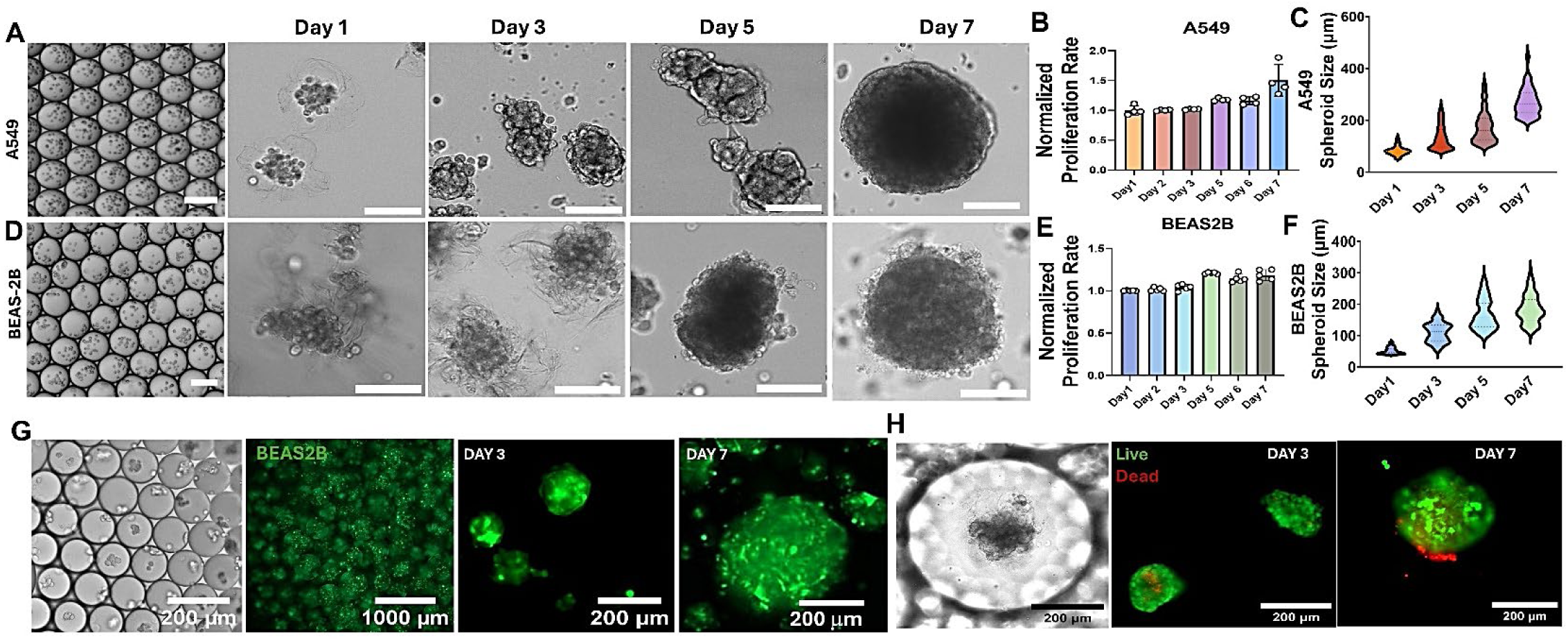
High throughput 3D cell culture using collagen microbeads. **(A)** Representative images on 3D culture of A549 lung tumor spheroids at different time points (Day 1, Day 3, and Day 7) using collagen microbeads. Scale bars represent 200 μm. **(B)** Quantitative analysis of fluorescence intensity (560/530 nm) for A549 spheroids from Day 1 to Day 7, indicating cell proliferation rate. **(C)** Average size of A549 lung tumor spheroids from Day 1 to Day 7, showing growth over time. **(D)** Representative images on 3D culture of BEAS-2B human epithelial alveolar spheroids at different time points (Day 1, Day 3, and Day 7) using collagen microbeads. Scale bars represent 200 μm. **(E)** Quantitative analysis of fluorescence intensity (560/530 nm) for BEAS-2B spheroids from Day 1 to Day 7, indicating cell proliferation rate. **(F)** Average size of BEAS-2B epithelial alveolar spheroids from Day 1 to Day 7, showing growth over time. Scale bars represent 200 μm. **(G)** Fluorescence images of BEAS-2B spheroids stained with Calcein AM (green, live cells) and Ethidium Homodimer-1 (red, dead cells) at Day 3 and Day 7. **(H)** Quantitative analysis of fluorescence intensity (560/530 nm) for A549 lung tumor spheroids from Day 1, Day 3 to Day 7, indicating cell viability.

### 3.4 3D manufacturing of extracellular vesicles

EVs have been recognized in the field for playing a crucial role in intracellular communication and various physiological processes[25, 38], which has been emerging in developing diagnostics and therapeutics. The biomimetic tissue microenvironment enabled by 3D cell cultures could promote cell– cell and cell–matrix interactions, in turn, stimulating EV production[39]. Importantly, 3D cell cultures have demonstrated the ability to enhance both the yield and therapeutic quality of EVs. In this study, we investigated the potential of collagen microbeads for sustained EV production as a potential drug delivery vehicle. We initially assessed the growth of C212 murine myoblasts in a concentration of 10×10^6^ in1 mg/mL collagen solution to form numerous cell-encapsulated collagen microbioreactors approximately 300 μm each in size (~141 cells per microbioreactor). We sustained the growth for over 30-day culture within the collagen microbioreactors (Figure 7A). We monitored EV production in 7 days using exosome-depleted media for culture media exchanges every two days. Myoblast cells exhibited an extended morphology with a network-like structure and filamentous actin cytoskeleton characteristics. Within the microbead, the collagen network was preserved clearly surrounded with cell clusters, which indicated ECM formation. On day 14, the media was collected, and extracellular vesicles were isolated using sucrose cushion ultracentrifugation method and characterized in terms of size, concentration, zeta potential, protein content and RNA content. Nanoparticle tracking analysis (NTA) revealed a mean size of ~170 nm particles in concentration of ~1×10^10^ particles/mL, and a zeta potential of ~-32 mV (Supporting Information Figure S4). As determined by micro bicinchoninic acid assay (MicroBCA), the average total EV protein concentration of 372 ng/mL was obtained. The RiboGreen RNA assay was also used to quantify total EV RNAs with an average concentration of 79 ng/µL. These results highlight the capability of collagen microbeads to support myoblast culture with sustained EV production, making them a potential tool for therapeutic applications.

**Figure 7.**
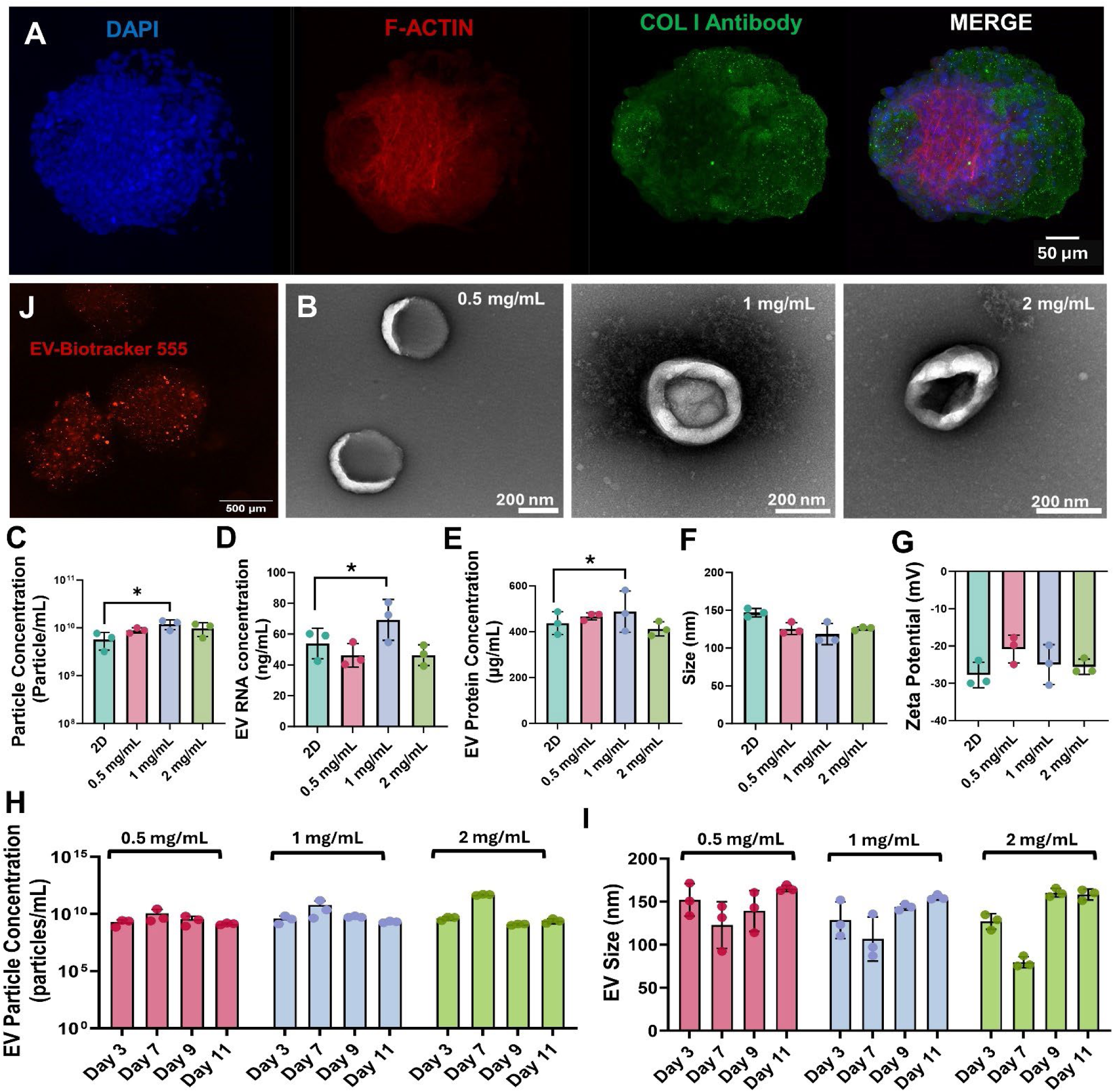
Collagen microbead 3D culture for extracellular vesicle production. **(A)** Immunofluorescence staining of C2C12 murine myoblasts cultured within a collagen microbioreactor for 14 days. Nuclei are stained with DAPI (blue), F-actin with phalloidin (red), and collagen type I antibody (green). The merged image shows viable culture and organization of cells within the collagen matrix. **(B)** TEM images showing the isolated EVs with typical morphology and size from 3D microbead culture in different collage concentrations (0.5 mg/mL, 1 mg/mL, and 2 mg/mL). **(C)** Quantification of EV particle concentration, **(D)** total EV RNAs, **(E)** total EV proteins, **(F)** Size, and **(G)** Zeta potential, from microbeads cultured in collagen concentrations of 0.5 mg/mL, 1 mg/mL, and 2 mg/mL, with significantly higher EV yield observed in 1 mg/mL, compared to 2D controls (p<0.05). **(H)** Assessment of EV particle concentration over cultures on days 3, 7, 9, and 11, showing sustained EV production across the duration of culture for each collagen concentration. **(I)** Assessment of EV size over cultures on days 3, 7, 9, and 11 for 0.5 mg/mL, 1 mg/mL, and 2 mg/mL collagen concentrations. **(J)** Fluorescently stained EVs within the microbeads showing retention of EVs uniformly distributed within the collagen matrix.

We then explored the potential of our microbioreactor system to enhance the production of EVs in comparison to gold-standard monolayer 2D culture[25]. The effect of different collagen concentrations in microbeads were assessed by suspending 5×10^6^ C2C12 murine myoblast cells in 0.5 mg/mL, 1 mg/mL, 2 mg/mL solution to form numerous collagen microbioreactors in size of approximately 300 μm (~70 cells per microbioreactor). At day 3 of culture, exosome depleted cell media was collected and the collagen microbioreactor cultures were digested using collagenase to release EVs inside. EVs were purified to image using Transmission Electron Microscopy, which confirmed consistent and characteristic EV morphology under different culture conditions (Figure 7B). Our findings suggest that at a concentration of 1mg/mL, C2C12 myoblasts secreted significantly more particles compared to the conventional 2D culture (p<0.05) (Figure 7C), as well as the higher content of total proteins (Figure 7D) and total RNAs (Figure 7E). Regarding EV size(Figure 7F) and zeta potential (Figure 7G), there was a slight decrease across the different collagen concentrations, compared to 2D culture. This 3D collagen microbead maintains cell viability and supports a network-like ECM microenvironment, indicating the healthy cell growth to preserve consistent EV production in good quality, in terms of size, protein, and RNA content, as well as zeta potential. We next repeated the encapsulation of 5×10^6^ cells/mL C2C12 myoblasts in 0.5 mg/mL, 1 mg/mL, and 2 mg/mL collagen microbioreactors to explore continuous and sustained EV release without disrupting the collagen matrix, establishing a self-contained EV release system. EVs were collected on days 3, 7, 9, and 11. EV Particle concentration analyses revealed a sustained EV production throughout the entire culture period (Figure 7H). The EV size remained average between 150-180 nm over time (Figure I). Overall findings suggest that developed collagen microbioreactors not only support enhanced EV yield and quality, but also ensure consistency and stability in EV production across long period, underscoring the platform’s robustness as a continuous EV production system.

Next, we tested the EV encapsulation and retention in collagen microbeads which can be used for in vivo controlled release and drug delivery. We collected EVs derived from gastrocnemius mouse muscle and performed density gradient ultracentrifugation for isolation. Afterwards, the purified EVs were stained with an orange cytoplasmic membrane dye and resuspended in 1 ml of PBS. To remove the free dye, we used ultracentrifugation with a 50% sucrose bed, which is confirmed for removing free dyes in the supporting Figure s5. A concentration of 10^12^ stained EVs were encapsulated in 1 mg/mL collagen type I microbeads (~600 μm). The free dye encapsulation was served as the control group. After ultracentrifugation, the fluorescence from the free dye group disappeared from microbeads. In contrast, EV group showed a stronger fluorescence imaging in Figure 7J and Figure s5, indicating an effective EV retention for potential application in delivery and controlled release in specific tissue sites.

## 4. Conclusions

In this manuscript, we utilize the collagen self-assembly under variable initial concentration and the change of temperature, which leads to a precise controlled collagen microstructure as the extracellular matrix. Such precisely controlled extracellular matrix environment allows for advanced applications in developing biomimetic models with physiological relevance. The most common collagen densities used in the field range from 0.5-5 mg/mL[40, 41], which is consistent with our described collagen microbeads using a concentration ranging from 0.25 to 3 mg/mL. It has been recognized that architecture of collagen networks seems to be linked to the conditions of polymerization. For instance, Jansen, et al. reported that at temperatures of 30°C and higher, the networks displayed characteristics of density, isotropy, and uniformity[42]. Drawing from the findings presented in this study, the mechanical forces exerted in the formation of the droplet playing a significant role in the formation of collagen networks and their spatial distribution within spherical geometry. The flow field corresponds to a Poiseuille-like profile where the velocity increases linearly from the wall to the center of the microchannel. Such dynamic force could result in collagen networks, lending further depth into generated spheroidal models. The captured SEM images in Figure 3 also showcased the tightly woven collagen network at concentration of 1mg/mL. In the context of individual micro-ECM generated with collagen, a matrix of 1 to 3 mg/mL is optimal for the growth and migration from tumor cells. Although many studies have suggested the use of spheroid for drug discovery and development, practical implementation has been constrained by challenges such as limited generational yield and lack of uniformity. Our approach offered an advanced solution for precise and high throughput production of collagen microbeads as the individual spheroid building blocks.

In the application of 3D culture for EV harvesting, collagen microbeads can serve as non-destructive, self-contained 3D culture vessels that enable sustained EV release. Currently, the main drawbacks of scaling EV production using 3D culture systems include inconsistencies in batch-to-batch variations, limited scalability, and the lack of non-destructive 3D scaffolds for sustainable EV manufacturing[25, 38]. Our platform addressed these challenges by producing up to 10,000 microbeads per minute, in geometries as small as 100 µm, with the capacity to encapsulate variable cell densities, which ensures both uniformity and high throughput, as well as consistent and sustainable EV collection in 3D cultures.

In this manuscript, we have demonstrated the successful fabrication and wide applicability of collagen microbeads. The production of collagen microbeads is facile and fast, which is suited for large scale 3D bio-ink production. We showcased the use of collagen type I microbeads at a concentration of 1mg/ml supplemented into an alginate/gelatin bioink, which maintains the mechanical properties of the bulk hydrogel while still containing large amount at a ratio of 1:1 of collagen type I with beneficial properties. The spheroidal conformations in the collagen microbeads could serve as functional microtissue within the printed constructs. Conventional 3D live cell printing is challenging and often lacks a particular cellular organization, due to the mechanical forces during hydrogel extrusion, coupled with viscosity changes like shear thinning or thickening, which can pose cell detachment, membrane disruptions and apoptosis. Our collagen microbead approach overcomes such challenges by providing formed spheroids as the intricate tissue architecture and enabling cell migration and distribution found in *in vivo* models. Herein, our approach opens the possibility of creating more realistic and physiologically relevant tissue models, in turn, leading to more accurate predictions of drug responses, deeper insights into disease progression, and more successful approaches to tissue repair and replacement.

## Acknowledgements

This project was supported by NIH NIGMS MIRA award 1R35GM133794 to Dr. Mei He. The authors would like to thank Anton Paar for use of their rheometer through the Anton Paar VIP research program. Authors would also like to acknowledge Pei Zhuang Ph.D. and Yi-Hua Chiang for their contributions in the development of this manuscript.

## Data statement

Research data is available upon request with proper citation.

## Author contributions

Conceptualization: Samantha Ali, Mei He

Data curation: Samantha Ali, Fabiana Mastantuono, Andrea Orozco-Torres, Niloy Barua

Funding acquisition and Project adminstration: Mei He

Resources: Xin Zhou, Lizi Wu Writing – original draft: Samantha Ali

Writing – review and editing: Samantha Ali, Mei He

