## Supplementary Materials for "High-throughput Generation of Collagen Microbeads for 3D Cell Culture and Extracellular Vesicle Production"

### SUPPORTING INFORMATION:

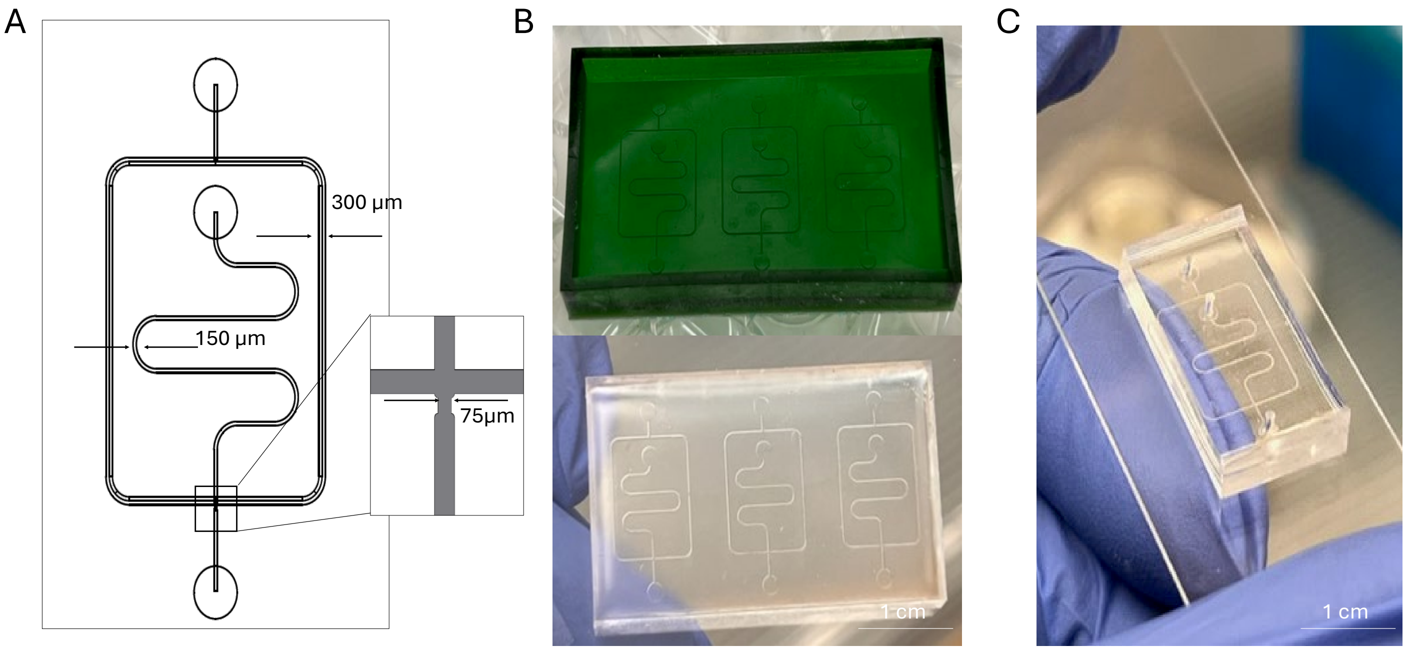


**Figure S1**. **A)** Schematic diagram of the microfluidic device design, illustrating the channel layout and the dimensions of the microchannels, with a cross-sectional view showing a channel width of 75 μm. **B)** 3D printed resin master mold used for the fabrication of microfluidic devices. **C)** Polydimethylsiloxane (PDMS) replica of the microfluidic device, which was cast from the master mold clearly showing the channels. **D)** Completed microfluidic device after plasma bonding of PDMS layer to a glass slide.

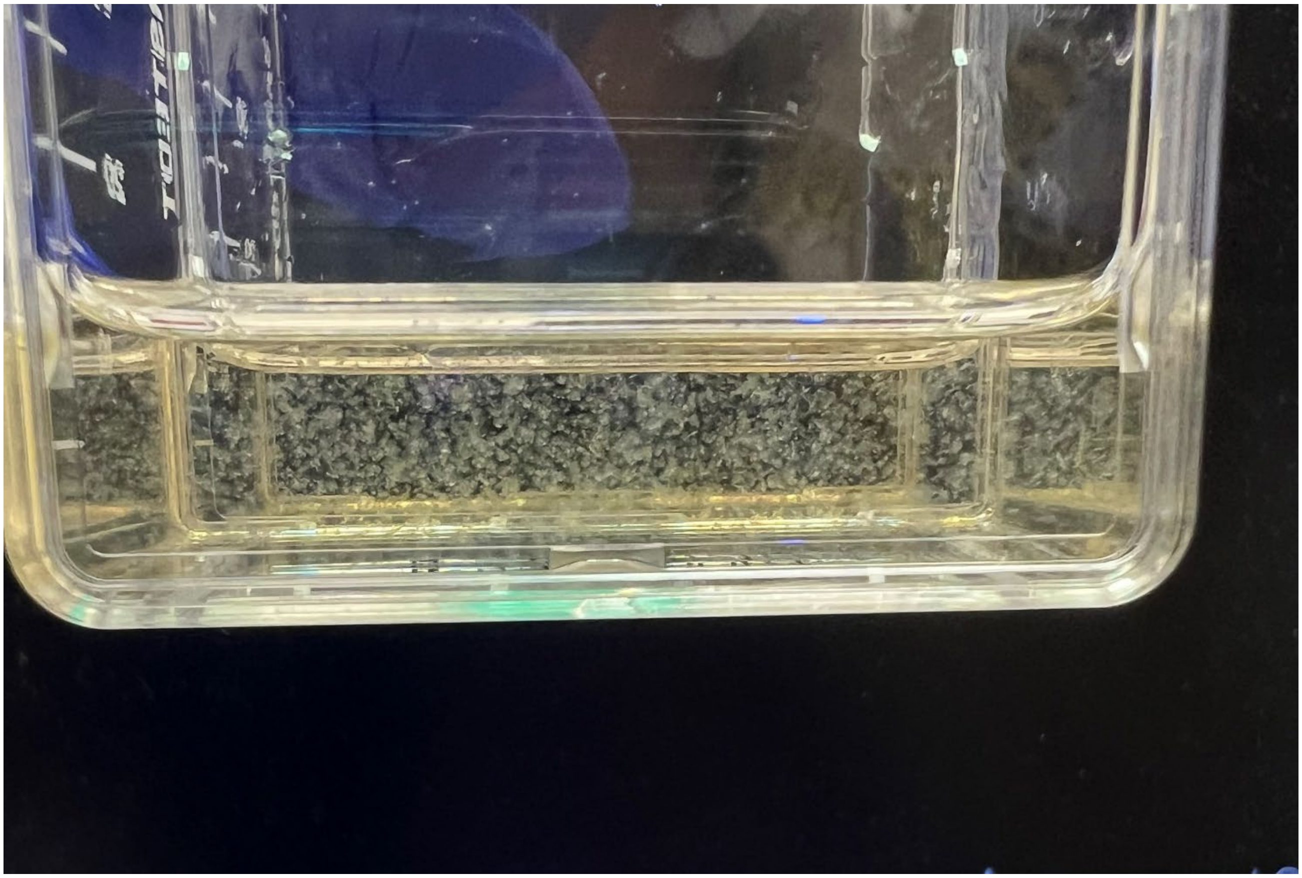

**Figure S2**. Collagen Microbeads Suspended in culture medium.

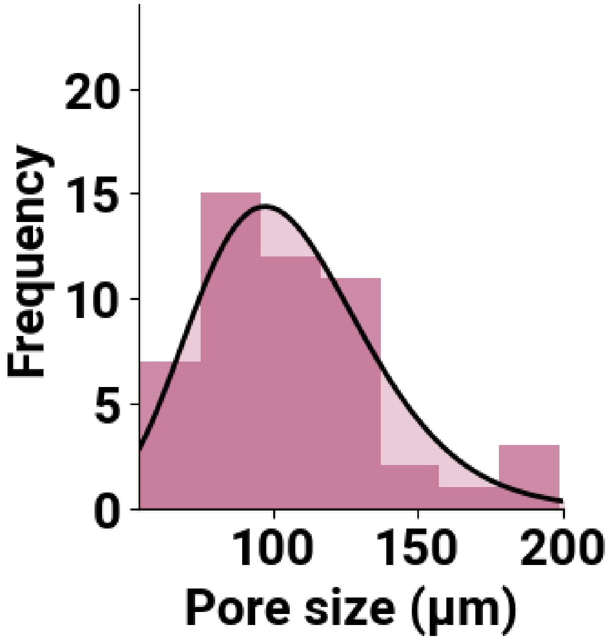


**Figure S3**. Pore size frequency distribution of honeycomb-like structures found in AG collagen microbead bio-ink.

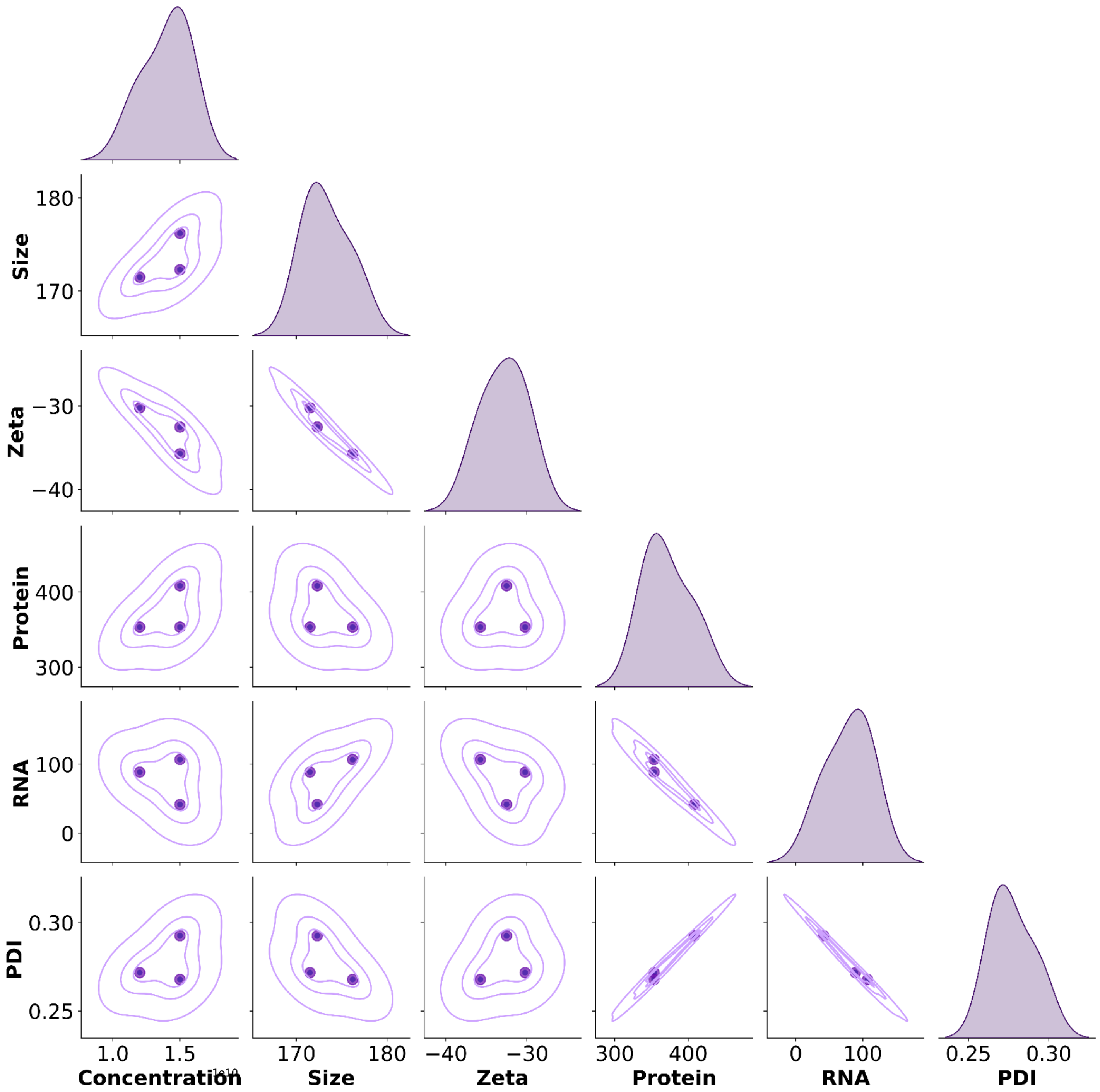


**Figure S4**. Pairwise comparison plots showing the relationships between various properties of EVs isolated from C2C12 murine myoblasts cultured in a 3D collagen microbioreactor. The analyzed properties include EV concentration (particles/mL), size (nm), zeta potential (mV), total protein content (μg), total RNA content (ng), and polydispersity index (PDI).

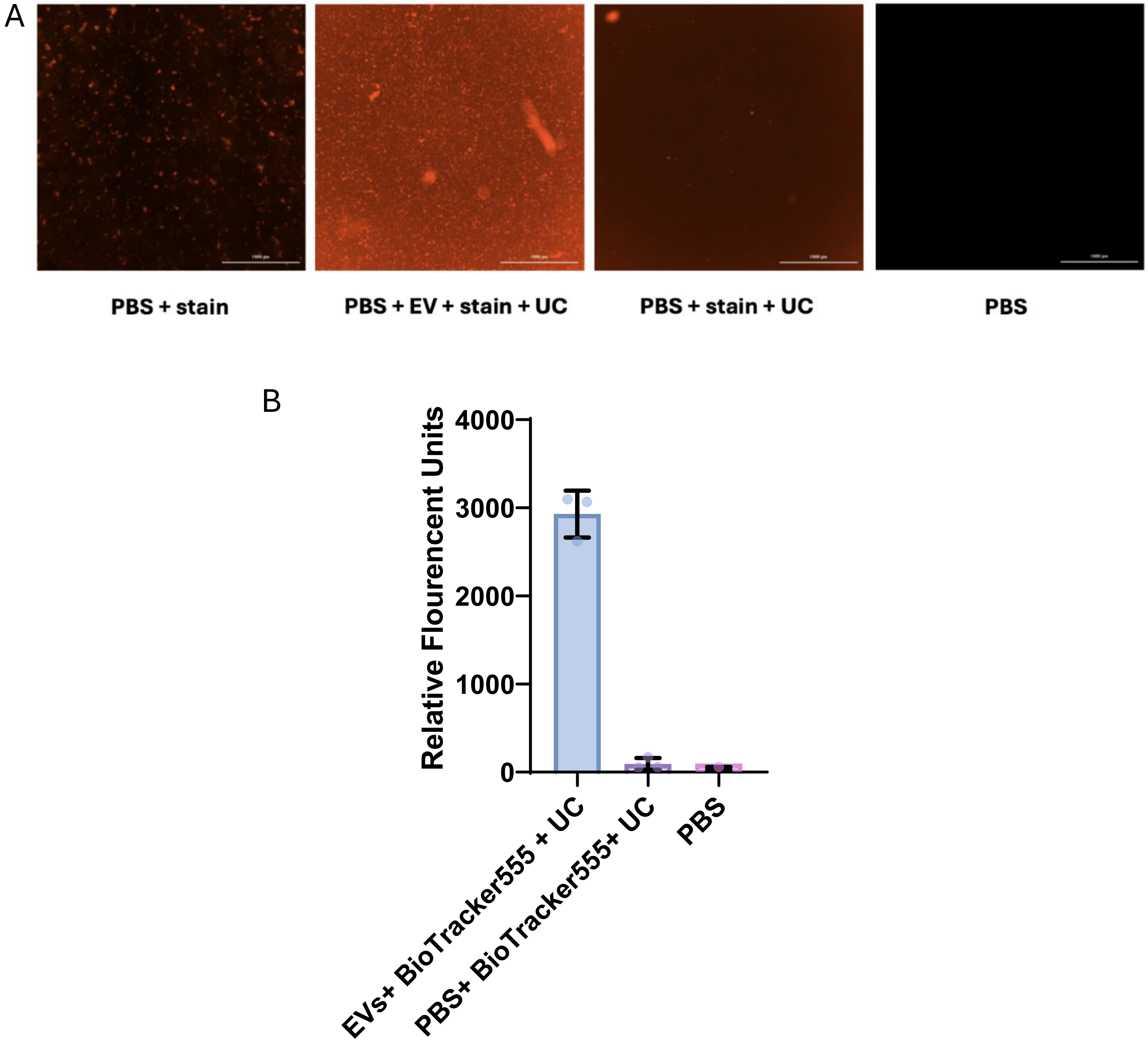


**Figure S5**. **A)** Representative fluorescence microscopy images of different control samples stained with cytoplasmic dye. The samples include (from left to right): Phosphate-buffered saline (PBS) with the cytoplasmic dye, PBS with extracellular vesicles (EVs) stained with the cytoplasmic dye and subjected to ultracentrifugation (UC).PBS with the cytoplasmic dye subjected to UC. Unstained PBS as a negative control. **B)** Quantification of measured fluorescence intensity for each sample. The fluorescence intensity values are presented as mean ± standard deviation (SD).
